# Sexual and temporal variation in New Zealand bellbird song repertoires

**DOI:** 10.1101/2021.09.29.462458

**Authors:** Michelle M. Roper, Wesley H. Webb, Yukio Fukuzawa, Christine Evans, Aaron M.T. Harmer, Dianne H. Brunton

## Abstract

How song repertoires vary within species and change over time is well studied in male songbirds. However, variation in female song repertoires remains largely unstudied despite female song being much more common and complex than once assumed. We investigated the song syllable repertoire of the New Zealand bellbird (*Anthornis melanura*), a species where both sexes have complex but sexually dimorphic song. We compared songs at individual and population levels to investigate sex and temporal variation of syllable repertoires. We detected 96 syllable types in the population over four years, of which 58% were unique to males, 32% unique to females and 9% were shared between the sexes. The population syllable repertoire of both sexes changed substantially across years with similar turnover rates (Jaccard’s similarity coefficients; female 52.9–69.0%; male 58.6–73.7%). Furthermore, many syllable types, unique to each sex, varied in prevalence within the population across years. The syllable repertoire sizes of individuals were higher for males than females (13-32, n = 7 and 6-16, n = 8, respectively). Although these sample sizes were low, the temporal variation in syllable prevalence and turnover for individuals were similar to patterns at the population level. Overall, male and female bellbirds exhibited similarities in temporal patterns of yearly repertoire composition, with rapid changes in syllable prevalence, but females had fewer syllable types than males. We suggest that these similarities and differences are consistent with male and female song repertoires being driven by similar but not identical selection pressures.

## Introduction

Until recently, female birdsong was considered a rarity rather than the norm. This was particularly apparent in the northern hemisphere, where in many species females either don’t sing or tend to sing less often than males (Arcese et al., 1988; Baptista et al., 1993). As a result, research focused on male song and the still prevailing paradigm that song repertoires are under direct selection by female choice and male-male competition (Catchpole, 1980). Contrasting with this paradigm, significant research has been undertaken showing that female song is the ancestral state for Oscines and is common in extant songbirds (Choe et al., 2021; Garamszegi et al., 2007; Odom et al., 2014; Riebel, 2003; Webb et al., 2016). However, quantitative studies of female song repertoires are still rare and there is a call for more integration of female song in birdsong research to provide a platform for better understanding female song function and evolution (Odom and Benedict, 2018; Riebel et al., 2005; Riebel et al., 2019). The prevalence of female song in species varies from no song, such as the zebra finch (*Taeniopygia guttata*; Zann, 1996), to species where females have comparable song repertoires to males, such as streak-backed orioles (*Icterus pustulatus*; Hall et al., 2010). This variation between species highlights the need to describe female song repertoires across a broader range of species from different geographic regions to explore what selection pressures are acting on the evolution of female song.

Song repertoires are under selection pressure from both inter- and intra-sexual interactions within species (Catchpole, 1980). Within a species, larger male song repertoires have been found to positively correlate with territory quality (Catchpole, 1986; Mcgregor et al., 1981) and size (Buchanan and Catchpole, 1997; Yasukawa et al., 1980), male parasite immunity (Buchanan et al., 1999), and reproductive success (Catchpole, 1986; Eens et al., 1991; Hasselquist et al., 1996; Lambrechts and Dhondt, 1986; Mcgregor et al., 1981). These correlations suggest that, for some species, song repertoires may be a trait used by a female to choose a mate. Alternatively, females may base mate choice on specific repertoire elements rather than just repertoire size alone, hence males may modify their repertoire to produce songs that are more attractive to females (Eriksen et al., 2011). In male canaries (*Serinus canaria*), particular syllables (an acoustic unit) and their spectral features have been referred to as ‘sexy syllables’; the production of these syllable types increases the number of copulation displays by females (Vallet et al., 1998; Vallet and Kreutzer, 1995). Song repertoire size may also be important in intra-sexual interactions; in experiments where male song is played in unoccupied territories, larger song repertoires can deter encroachment by neighbouring males more effectively than smaller repertoires (Krebs et al., 1978).

As in males, female song may have a role in both inter- and intra-sexual interactions (Riebel, 2003). For example, female red-winged blackbirds (*Agelaius phoeniceus*) have two song types; one produced in the presence of males, particularly their mate, and the other in the presence of females (Beletsky, 1983, 1985). Female song is also used differently in different mating and social systems, for example, cooperative breeders such as superb fairy-wrens (*Malurus cyaneus*) sing more towards unfamiliar females (Cooney and Cockburn, 1995). In contrast, socially monogamous breeders, such as New Zealand bellbirds (*Anthornis melanura)*, sing more in response to neighbouring females (Brunton et al., 2008). In species with more complex mating systems where females must compete for males, female song is used for mate attraction (Langmore and Davies, 1997; Langmore et al., 1996). Female song has also been linked to reproductive success, for example, higher singing rates towards territorial intruders (Cain et al., 2015), spontaneous female song and song complexity (Brunton et al., 2016) are positively correlated with the number of fledglings females’ produce. Female song may also function in within-pair communication (Rose et al., 2020; Sikora et al., 2021). These studies suggest that different aspects of female song repertoires may be driven by either (or both) intra-sexual and inter-sexual selection pressures.

Even fewer studies are available that shed light on how female song repertoires are learnt, develop and change over time. Furthermore, most of this research has been done on species where males and females significantly overlap in the song components of their repertoires (shared syllable types). These studies demonstrate that developmental patterns can be similar but not identical to those seen for male conspecifics (Roper et al., 2018). Common developmental patterns include increasing or decreasing song repertoires with age, early attrition of syllables, and varying stability of syllable longevity and prevalence throughout an individual’s lifetime. For example, female alpine accentors (*Prunella collaris*) increase their syllable repertoire size with age, particularly between their first and second year of age (Langmore et al., 1996). In contrast, the syllable repertoires of female blue-capped cordon-blues (*Uraeginthus cyanocephalus*) have higher levels of selective early attrition (i.e., discarding syllables from their repertoire that were learnt prior to song crystallisation) than males (Lobato et al., 2015). Hence while both sexes have a similar syllable repertoire size as juveniles during development, adult female cordon-blues have a significantly smaller syllable repertoire size than males (Lobato et al., 2015). Furthermore, in slate-coloured boubous (*Laniarius funebris*), juvenile females occasionally innovate syllable types during song development, whereas males do not (Wickler and Sonnenschein, 1989). Studies of song repertoires of female European starlings have been inconsistent; Pavlova, Pinxten, & Eens (2010) found female song repertoires decrease with age, whereas other studies have shown female starlings alter and increase their repertoire size with age by adding new and deleting old phrase types (a sequence of syllables) as males do (Pavlova et al., 2010). This research indicates that in some species females can change their song repertoire over time, but few studies (e.g. Graham et al., 2021) have examined species with a high degree of sexual dimorphism in their song repertoires.

Here, we investigated sexual and yearly temporal variation in the song repertoires of the New Zealand bellbird (*Anthornis melanura*; hereafter bellbird); a species that is exceptional compared to previously studied species with female song, as their song is highly sexually dimorphic (Brunton and Li, 2006). Brunton and Li (2006) found that, on average, females have a smaller repertoire size of 1.9 song types versus 5.4 for males, but this could be an underestimate due to the sampling methods used (the range was from one to six song types for individual females). At the population level, the authors detected 36 syllable types: 17 unique to males, 12 unique to females, and seven syllable types shared by the sexes. Females tend to sing shorter and more variable song bouts than males, but despite both sexes having high year-round singing rates, females sing more than males in the breeding season (Brunton and Li, 2006). Both sexes develop their songs in a comparable timeframe (Roper et al., 2018) and possible functions of female song include territory defence, polygyny prevention and sexual selection (Brunton et al., 2008; Brunton et al., 2016).

The aim of our study was to quantify the full syllable repertoire of male and female bellbirds on Tiritiri Matangi Island at both the population and individual level to compare the syllable repertoire of each sex, the degree of syllable sharing and determine the range of syllable types sung over time. We predicted that we would find more syllable types than Brunton and Li (2006) due to the longer duration of this study and more sampling per individual bird, but that the relative abundance of female to male syllable types and proportion of shared syllable types would be similar. We recorded bellbird song across four years and took both cross-sectional (population level) and longitudinal (individual level) approaches to answer two questions: 1) how does the syllable repertoire of each sex compare and 2) do syllable repertoires change at a similar rate for both sexes. Species where male and female song are largely distinct and complex provide an exciting opportunity to understand the evolution of female song.

## Methods

### Study site and data collection

We recorded bellbird songs for this study on Tiritiri Matangi Island. Tiritiri Matangi Island is a low-lying 220 ha island in the Hauraki Gulf, 4 km off the coast of the Whangaparaoa Peninsula and 25 km north of Auckland, New Zealand. We visited the island four days per week during the breeding season (August to January) and post-breeding season (March to May), between September 2012 and January 2016. We recorded bellbirds between the hours of 0800 to 1800 while searching for individually colour-banded birds and un-banded birds on territories. This study therefore excludes dawn chorus song, which has only been observed to be sung by males (Roper and Brunton, personal observation). Post-breeding season, bellbirds formed loose flocks near food resources (e.g., kanuka trees, *Kunzea ericoides*, for honeydew, or supplementary sugar water feeders; Roper, personal observation), where we targeted individuals for recording.

On a regular basis, we colour banded individual bellbirds to allow individual repertoires of adult males and females to be recorded. We captured the adults using mist nets on bellbird territories or at specially modified cages containing sugar-water feeders. The birds were banded with unique combinations of three colour bands, and one metal band. All capture, handling and banding protocols were conducted under permits from the New Zealand Department of Conservation (20666-FAU, 34833-FAU, 2008/33) and the Massey University Animal Ethics Committee (12/32, 15/21). We avoided capturing near nests when we knew a female was incubating.

### Audio recording and acoustic processing

We recorded bellbird song using a directional shotgun microphone (Sennheiser ME66, Sennheiser, Germany) and a portable solid state recorder (Marantz Professional PMD 661, Marantz Professional, Cumberland, RI, U.S.A.) with 24-bit sampling precision and 48 kHz sampling rate. Recordings were made of colour-banded individuals that were banded prior to and during the study and un-banded individuals. Recordings were made from the austral spring to autumn over the course of the study. The male-female ratio in the population was approximately equal based on previous surveys (Roper, 2012) and there is no evidence of sex-biased mortality (Baillie, 2011). However, there was a male bias in detecting individuals because males were more conspicuous than females, as males tended to dominate the supplementary sugar-water feeders (Roper, 2012). Total recording time was approximately 110 hrs with 2557 recordings (note that this is not an indicator of total field recording effort, as it does not include time spent following individuals waiting for singing to occur).

Recordings were processed in Raven Pro 1.5 (Cornell Lab of Ornithology, Ithaca, NY) with standardised settings (a bandpass filter of 500 Hz to 22 000 Hz, size 256 samples, time grid overlap 50%, and normalised by amplifying to fill the waveform window). Songs or bouts of singing were selected (hereafter ‘song selection’) from each raw recording. We created Excel (Microsoft Corporation) databases to enter metadata for each raw recording and song selection. The highest quality song selections were then imported into the open-source software Luscinia (Lachlan, 2007).

We chose syllables as the acoustic unit for measuring repertoire size to compare between the sexes, as males and females sing different song types and the structure of their songs differ (Brunton and Li, 2006). We defined syllables as consisting of one or more elements (i.e., note; a single sound trace on spectrogram) with less than a 15 ms interval between elements or the elements were consistently repeated together within a song type (Kershenbaum et al., 2016; Marler and Isaac, 1961). See Figure 1 for examples of male and female bellbird syllable types. Elements and syllables were selected from spectrograms (FFT algorithm) of song samples using Luscinia (Lachlan, 2007). We used the following settings for viewing spectrograms in Luscinia, as recommended by R. Lachlan (personal communication): max. frequency 12 000 Hz; frame length 10.667 ms; spectrograph points 470; spect. overlap 90.63%; deverberation was set to 100% to reduce echo; dynamic range was altered between 30 and 60 to obtain the best estimation of fundamental frequency when selecting elements; high pass threshold 300 Hz.

**Figure 1.**
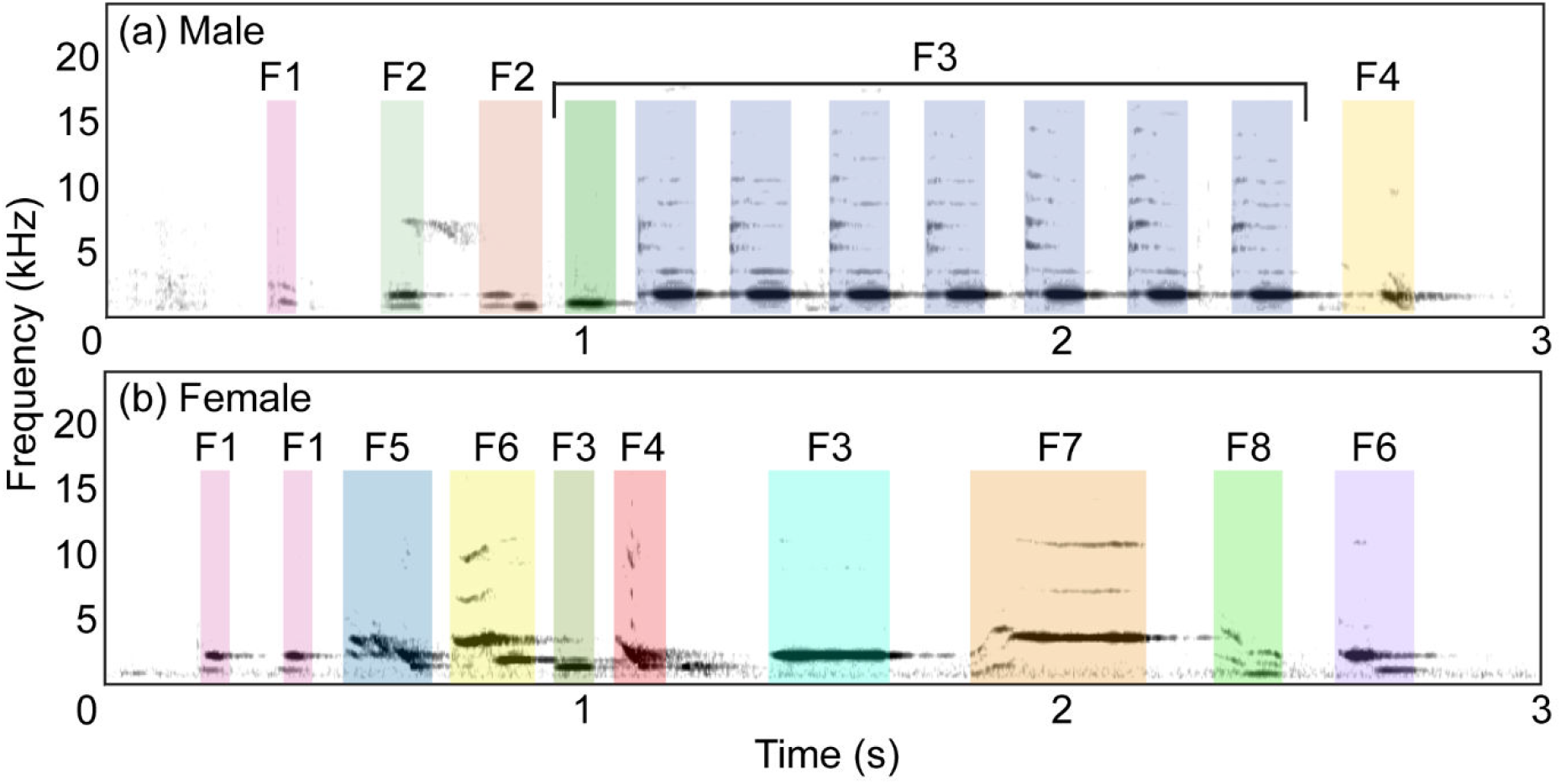
Examples of bellbird syllable types (different syllable types are represented by a different coloured box) for (a) male and (b) female bellbirds with syllable family types labelled with the following codes: F1 = stutter, F2 = waah, F3 = pipe, F4 = chump, F5 = down transition, F6 = step-down, F7 = flat-squeak, F8 = down-squeak.

The syllables selected in Luscinia were then imported into the open-source database program Koe (www.koe.io.ac.nz; Fukuzawa et al., 2020) where syllables were systematically labelled into categories. Syllable types were categorised by eye and ear based on a range of features including shape of elements, fundamental frequency, frequency modulation, harmonics and duration. One experienced observer then used additional syllable ordering information within the song types (i.e., where a syllable type was sung within a song) to ensure consistent labelling of the syllables and agreement was made with an additional experienced observer. For example, if two very similar syllable types were sung with the same syllable order in each song, they were merged together as one syllable type. To verify that the labelling was consistent, machine learning algorithms (supervised and unsupervised) were used to assess how well the syllable labels clustered using the algorithm t-SNE (t-distributed Stochastic Neighbour Embedding; van der Maaten and Hinton, 2008) and determined the accuracy rate using various machine learning algorithms (see Supplementary Methods 1 for additional methods). The syllable labels showed a high degree of clustering (Supplementary Figure 1) and syllables were labelled with 96.6 ± 1.0% accuracy (Supplementary Table 1). Additionally, syllables were grouped into broader ‘syllable family’ groups, where syllable types with similar spectral characteristics were given a family label (e.g., pure tone syllables were in the family ‘pipe’; see Figure 1 for examples of syllable family types and see Supplementary Table 2 for examples from all syllable family types).

### Population syllable repertoire size

Using the Koe database, we summarised the number of syllable types and syllable families for the entire study period and between years. A year of recording consisted of both the breeding season months and post-breeding season months, and the four years of recording were numbered consecutively (Year 1 = October 2012 to May 2013, Year 2 = September 2013 to April 2014, Year 3 = September 2014 to March 2015, and Year 4 = September 2015 to January 2016). We used simple enumeration to estimate the number of syllable types by plotting cumulative curves of observed syllable types versus number of song selections for each sex across all years and each year with package ggplot2 (Wickham, 2016) in R Studio (R Core Team, 2015). This was done to determine whether the entire population syllable repertoire was captured, determined by whether the cumulative curve approached an asymptote. For syllable types, we calculated the total number of types in the population, number of types sung by each sex and how many types were shared by the sexes, across all years of recording and separately for each year.

### Variation in population syllable repertoire over time

To determine how the population repertoire changed over time (i.e., yearly turnover rate), we compared the syllable types present each year using Jaccard’s similarity coefficient (MacDougall-Shackleton et al., 2009; Podos et al., 1992; Tracy and Baker, 1999), their presence and absence between years, and relative abundance in the population. To measure the similarity of syllable types across years separately for males and female, we calculated Jaccard’s coefficient (*S*_*j*_) following Tracy and Baker (1999) as:

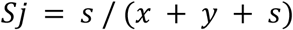

where *s* = number of syllable types shared by year X and year Y, *x* = number of syllable types present in year X but not in year Y, and *y* = number of syllable types present in year Y but not in year X. To compare the similarity across years between males and females, we used a two-sample t-test. The yearly totals of syllable additions, syllable deletions and shared syllable types was calculated to assess whether any differences in similarity were due to additions or deletions of syllable types from the repertoire or a combination. Since some syllable types may have been undetected in the repertoire due to rarity (and hence were counted as deletions), we calculated the relative abundance of each syllable type. For each syllable type, the total number of individuals singing that type was tallied and we calculated the proportion of individuals singing each type for each year. We used four categories of abundance based on the percentage of individuals singing each syllable type: rare <15%, uncommon 15–45%, common 46–75%, and abundant >75%. To determine if any syllable changes were occurring at small scale (i.e., within syllable family type) or large/novel scale (i.e., new syllable family additions or old family deletions), we totalled the number of syllable types within each syllable family across each year.

### Individual syllable repertoire and variation over time

We used the same enumeration method as at the population level to calculate syllable repertoire size for individuals of each sex. We used a subset of individuals for which we had sufficient samples to estimate repertoire size (ten or more song selections). For each sex, we calculated the observed syllable repertoire size range and mean ± standard error (SE) for this subset of individuals and tested for differences between sexes with an un-paired two-sampled t-test. We also calculated the number of syllable types within each syllable family for this subset to compare the level of syllable diversity between individuals of each sex. We calculated the relative abundance of syllable types for individuals, using the same method as for the population level, to confirm the results found for syllable abundance when using all birds recorded in the population. To compare individual repertoire sizes between years, we selected individuals that had three or more song selections per year (we did not have the ten selections needed to approach an asymptote for observed repertoire size). As for the population, we calculated Jaccard’s similarity coefficient for the syllable types present between years and tallied the number of syllable additions and deletions.

## Results

### Population repertoire

A cumulative plot of the number of syllable types versus number of song selections approached an asymptote, showing the majority of the population syllable repertoire was captured (Figure 2). In total across the four years of the study, we identified 96 syllable types (Table 1). Of these, 58.3% of syllable types were unique to males, 32.3% were unique to females and the sexes shared 9.4% of syllable types (Table 1). The male population syllable repertoire was hence almost twice as large as the female population syllable repertoire. The syllable types were categorised into 20 syllable family types (Table 2). There were six families unique to males, one family unique to females, and 13 shared between the sexes.

**Figure 2.**
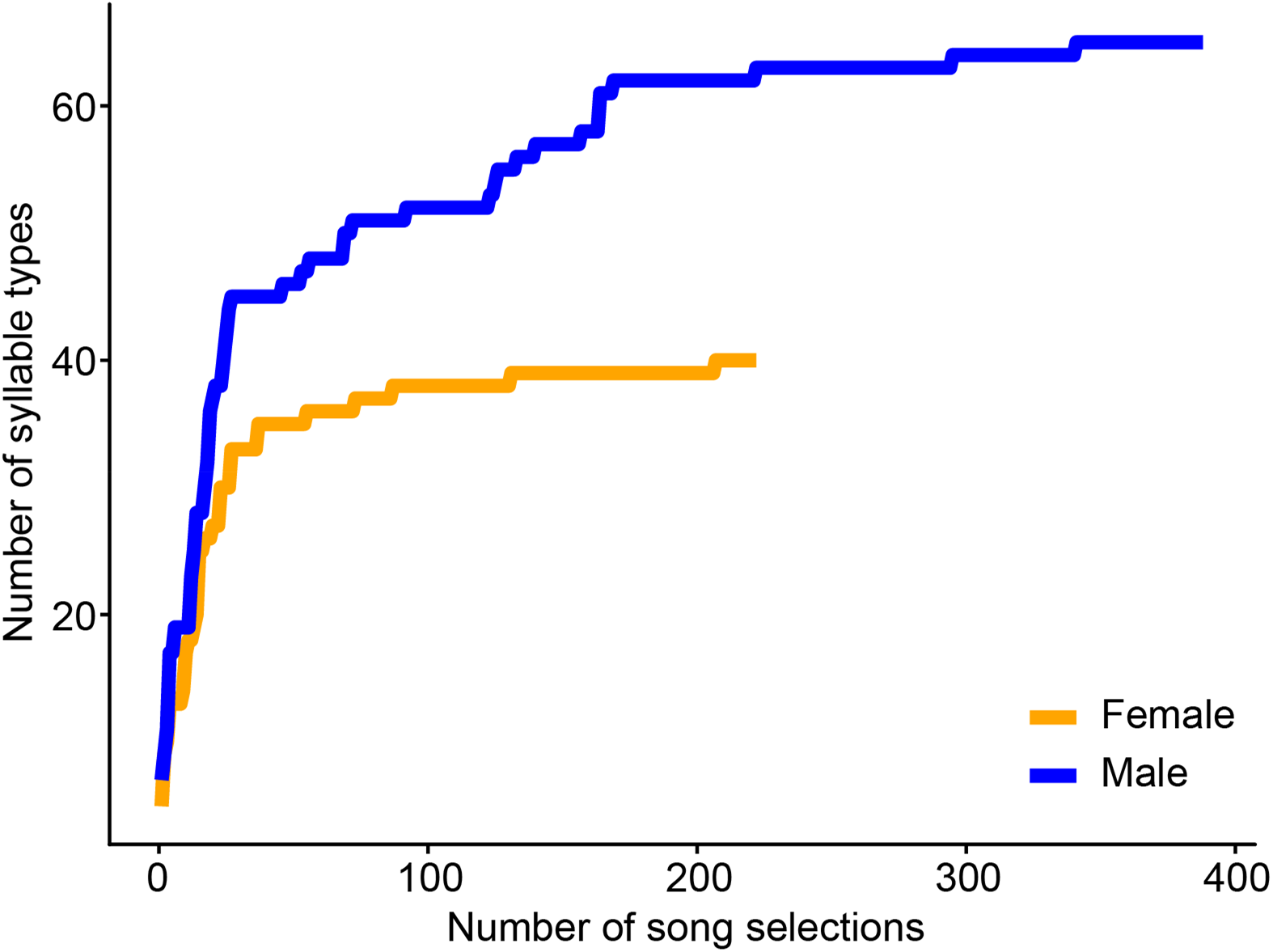
Cumulative plot of the number of syllable types versus number of song selections for male and female bellbirds on Tiritiri Matangi Island from year 1 (2012–2013) to year 4 (2015–2016).

**Table 1.** Number of syllable types across sexes and recording years (Year 1 = 2012–2013, Year 2 = 2013–2014, Year 3 = 2014–2015, Year 4 = 2015–2016).

|  | All years | Year 1* | Year 2 | Year 3 | Year 4 |
| --- | --- | --- | --- | --- | --- |
| Total | 96 | 60 | 72 | 74 | 71 |
| Male | 65 | 42 | 48 | 50 | 49 |
| (number of individuals) | (110) | (32) | (26) | (31) | (21) |
| Female | 40 | 21 | 31 | 29 | 28 |
| (number of individuals) | (62) | (10) | (18) | (19) | (18) |
| Shared | 9 | 3 | 7 | 5 | 6 |

\* Note that the female cumulative syllable count did not approach an asymptote in this year.

**Table 2.** The number of syllable types present within each syllable family in the population repertoire of each sex for each recording year.

| Syllable family | Male |  |  |  | Female |  |  |  |
| --- | --- | --- | --- | --- | --- | --- | --- | --- |
|  | Year 1 | Year 2 | Year 3 | Year 4 | Year 1* | Year 2 | Year 3 | Year 4 |
| Alarmy | 2 | 1 | 1 | 1 |  |  |  | 1 |
| Chortle | 2 | 2 | 2 | 2 |  |  |  |  |
| Chump | 1 | 1 | 1 | 1 | 1 | 1 | 1 | 1 |
| Click | 1 | 1 | 1 | 1 |  | 1 | 1 | 1 |
| Cough | 1 | 1 | 1 | 1 |  |  |  |  |
| Down-squeak | 1 | 1 | 1 | 1 | 1 | 2 | 1 | 1 |
| Down-squeak pipe | 1 | 2 | 2 | 2 |  |  |  |  |
| Down transition |  |  |  |  | 4 | 5 | 4 | 4 |
| Flat-squeak | 1 | 1 | 3 | 1 | 1 | 1 | 1 | 1 |
| Peak-squeak | 1 | 2 | 2 | 2 | 1 | 1 | 2 | 2 |
| Pipe | 11 | 12 | 9 | 11 | 5 | 8 | 9 | 6 |
| Pipe down-squeak | 2 | 2 | 3 | 4 | 1 | 1 | 1 | 1 |
| Step-down | 2 | 2 | 2 | 2 | 3 | 3 | 4 | 4 |
| Step-up | 1 | 1 | 2 | 1 |  |  |  |  |
| Stutter | 2 | 2 | 2 | 2 | 4 | 4 | 4 | 6 |
| Stutter up-squeak | 1 | 1 | 1 | 0 |  | 2 |  |  |
| Trill | 2 | 4 | 4 | 3 |  |  |  |  |
| Up-squeak | 1 | 2 | 1 | 1 |  | 1 |  |  |
| Waah | 8 | 9 | 11 | 12 |  | 1 | 1 |  |
| Warble | 1 | 1 | 1 | 1 |  |  |  |  |

\* Note that the female cumulative syllable count did not approach an asymptote in this year.

### Variation in population syllable repertoire over time

The cumulative plot showed the syllable repertoire approached an asymptote for each year except for females in year 1 (Supplementary Figure 2). The similarity coefficient (*S*_*j*_) for syllable types present between years fell within a narrow range from 52.9 to 73.7% (Table 3) with no significant difference between the sexes (*t* = -1.14, df = 9.91, *P* = 0.28). If the syllable repertoire changed due to deletions and additions over time, we expected that the further the years were apart (i.e., temporal distance), the less similar the syllable repertoire would be. However, we did not find a consistent trend. The average similarity between years with 1-year difference was 69.4% and 60.2%, 2-years difference 62.9% and 61.0%, and 3-years difference (not an average as only one sample) 62.5% and 69.0%, for males and females respectively. This suggests that syllable types may change in their relative abundance (i.e., become rare or more abundant) between years rather than being permanently deleted from the repertoire. Between most years (except for Year 1), the number of syllable types added and deleted were very similar (Table 3). The changes in syllable types between years were always within the existing family types (Table 2), except for females; although these new family types for females were existing male syllable family types rather than novel family types. For both males and females, the proportion of syllable types that were commonly or abundantly sang by individuals in the population was 14% or lower (Figure 3). Typically, the greatest proportion of syllable types were rare or uncommon among individuals (Figure 3), i.e., most syllable types were only sung by 45% or less of the individuals recorded.

**Table 3.** Syllable additions and deletions, number of syllable types shared and syllable repertoire similarity (Jaccard’s similarity coefficient; proportion) between years, compared at the population and individual level. See Supplementary Table 4 for the number of song selections for each individual per year.

| Comparison | Sex |  | Years | Added | Deleted | Shared | Similarity (%) |
| --- | --- | --- | --- | --- | --- | --- | --- |
| Population | Male |  | 1–2* | 13 | 7 | 35 | 62.5 |
|  |  |  | 2–3 | 9 | 7 | 41 | 71.9 |
|  |  |  | 3–4 | 7 | 8 | 42 | 73.7 |
|  |  |  | 1–3* | 16 | 8 | 34 | 58.6 |
|  |  |  | 2–4 | 10 | 9 | 39 | 67.2 |
|  |  |  | 1–4* | 14 | 7 | 35 | 62.5 |
|  | Female |  | 1–2* | 13 | 3 | 18 | 52.9 |
|  |  |  | 2–3 | 6 | 8 | 23 | 62.2 |
|  |  |  | 3–4 | 6 | 6 | 23 | 65.7 |
|  |  |  | 1–3* | 9 | 1 | 20 | 66.7 |
|  |  |  | 2–4 | 7 | 10 | 21 | 55.3 |
|  |  |  | 1–4* | 8 | 1 | 20 | 69.0 |
|  | Sex | Individual | Years | Added | Deleted | Shared | Similarity (%) |
| Individual | Male | 1 | 3–4 | 2 | 8 | 22 | 68.8 |
|  |  | 2 | 3–4 | 5 | 5 | 3 | 23.1 |
|  |  |  | 1–3 | 0 | 11 | 8 | 42.1 |
|  |  |  | 1–4 | 3 | 14 | 5 | 22.7 |
|  |  | 3 | 1–2 | 7 | 7 | 10 | 41.7 |
|  | Female | 1 | 3–4 | 0 | 2 | 13 | 86.7 |
|  |  | 2 | 1–2 | 2 | 6 | 3 | 27.3 |
|  |  |  | 2–3 | 0 | 1 | 4 | 80.0 |
|  |  |  | 1–3 | 1 | 6 | 3 | 30.0 |
|  |  | 3 | 3–4 | 0 | 0 | 8 | 100.0 |
|  |  | 4 | 3–4 | 0 | 3 | 8 | 72.7 |

\* Note that the female cumulative syllable count for the population did not approach an asymptote in this year.

**Figure 3.**
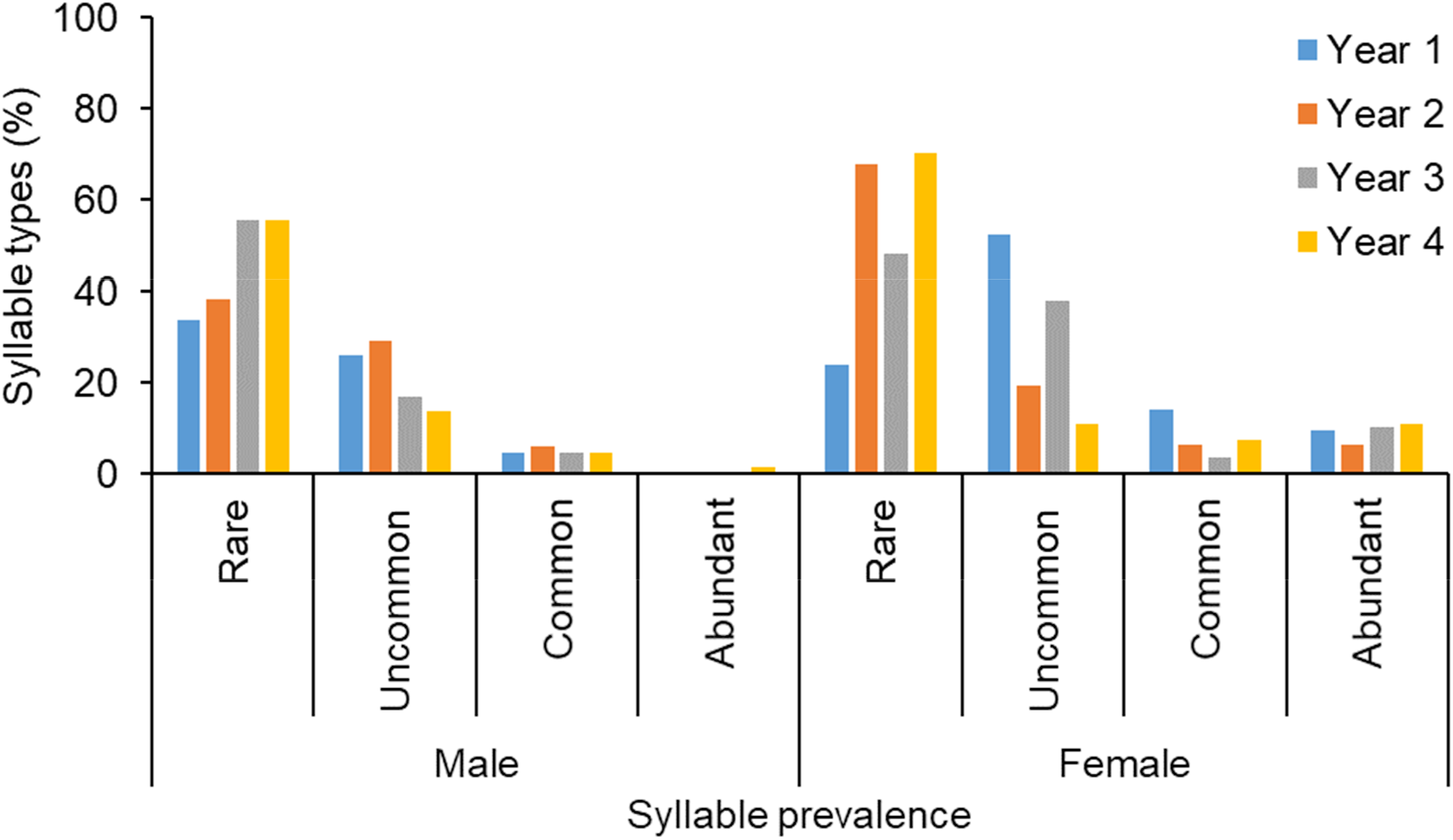
Prevalence of syllable types among the population repertoire compared against the proportion of individuals (rare <15%, uncommon 15–45%, common 46–75%, abundant >76%) singing each syllable type for each recording year (Year 1 = 2012–2013, Year 2 = 2013–2014, Year 3 = 2014–2015, Year 4 = 2015–2016). See Table 1 for the number of individuals within each year.

### Individual syllable repertoire and variation over time

Seven males and eight females had syllable repertoires that either approached or began to approach an asymptote (Figure 4). As the full repertoire size of each individual was potentially not captured, the mean observed repertoire size represents the theoretical minimum mean. The mean ± SE male syllable repertoire was 19.1 ± 2.5 and ranged from 13 to 32 syllable types (Supplementary Table 3). The mean ± SE female syllable repertoire size was significantly smaller than males at 10.8 ± 1.4 syllable types (*t* = -2.91, df = 9.26, *P* = 0.02) and ranged from six to 16 syllable types (Supplementary Table 3). There was variation in the number of syllable family types that individuals sang (Supplementary Table 3). Some syllable family types, however, were only sung by one individual. For example, only one male sang a syllable type in the family ‘stutter up-squeak’ and one female sang a syllable type in the family ‘waah’ (which is more commonly sung by males). As for all individuals at the population level, syllable type prevalence among individuals showed similar trends with most syllable types being sung by few individuals (rare and uncommon), and fewer syllable types sung by many individuals (common and abundant; Figure 5).

**Figure 4.**
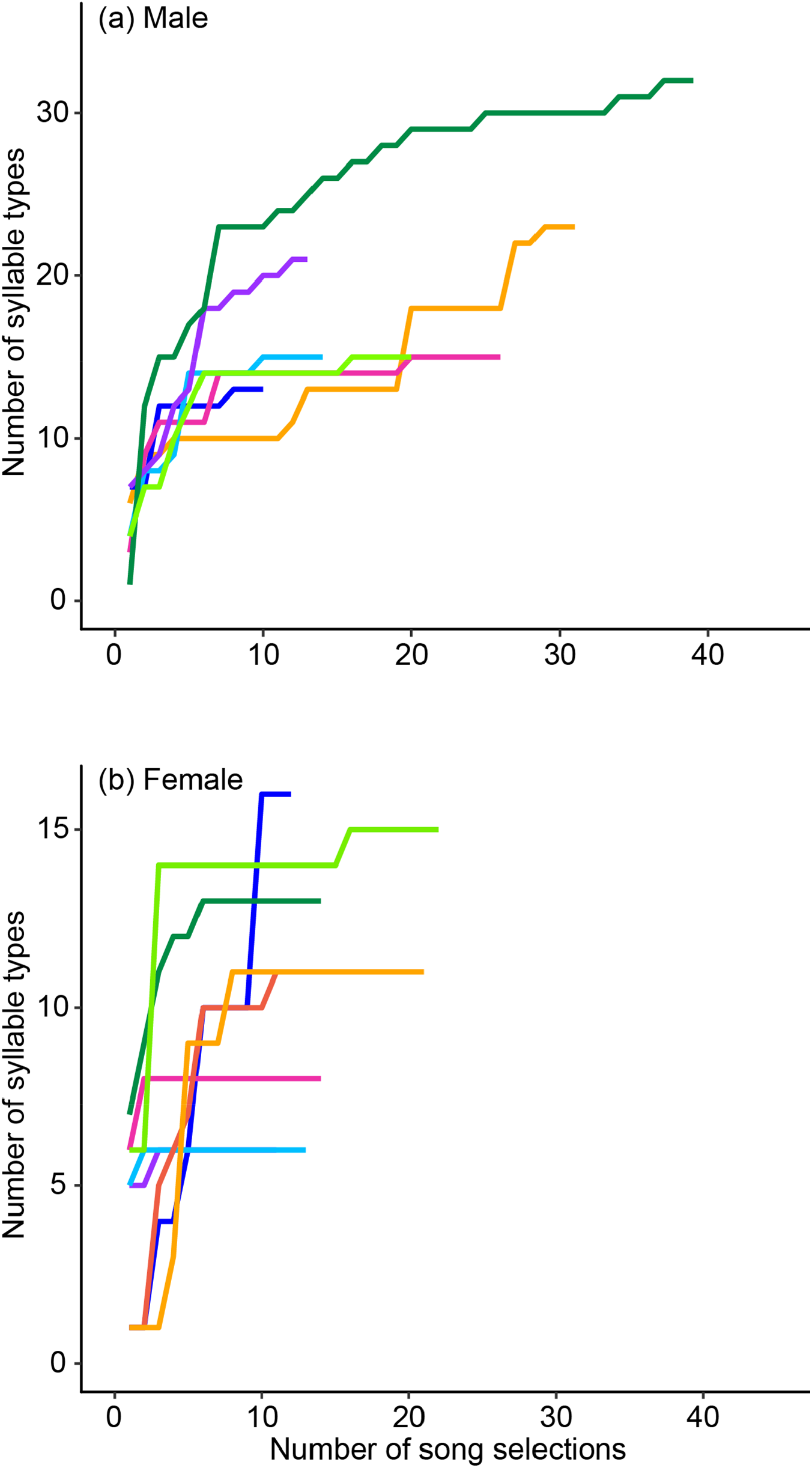
Cumulative plot of the number of syllable types versus number of song selections for individual (a) male and (b) female bellbirds (corresponding to the individuals in Supplementary Table 3) to estimate their total syllable repertoire size.

**Figure 5.**
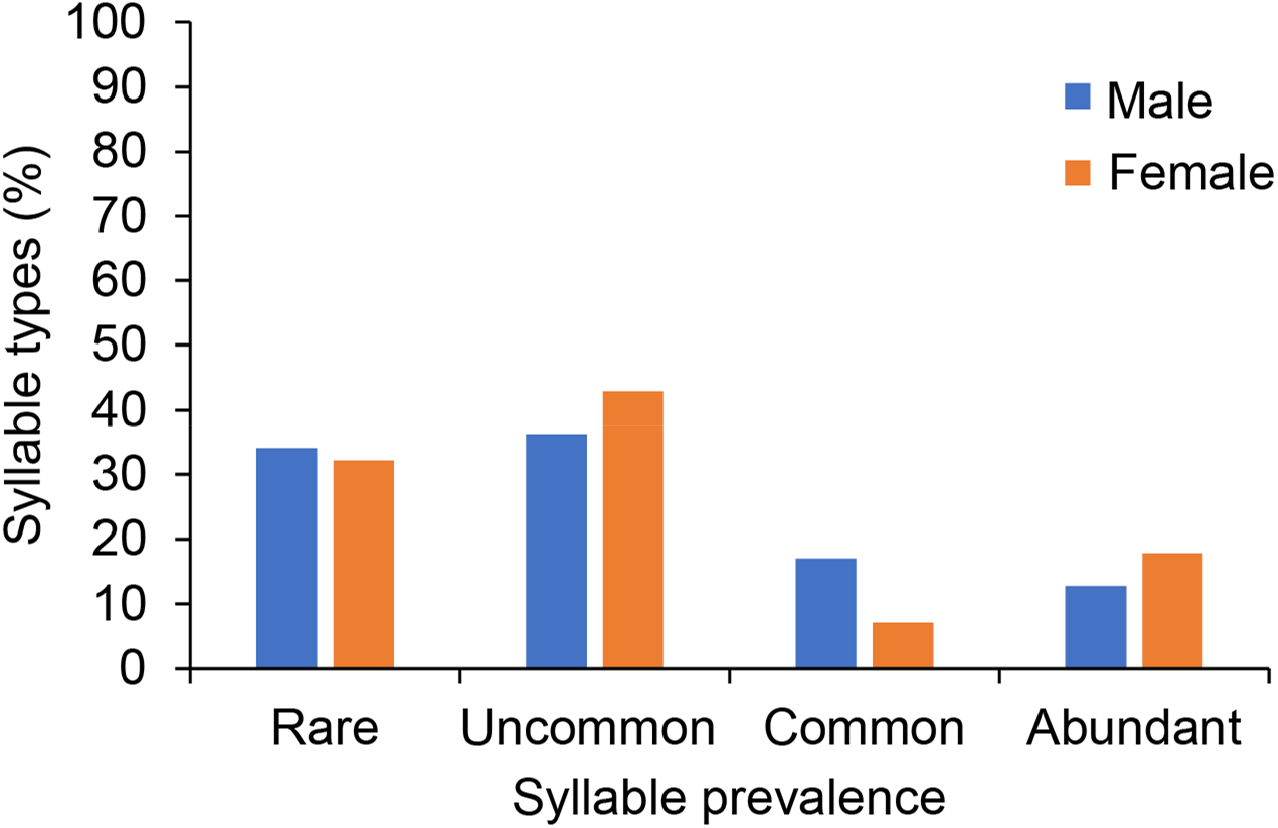
Syllable prevalence based on proportion of individuals (rare <15%, uncommon 15– 45%, common 46–75%, abundant >76%) singing each syllable type for the individuals where the full repertoire was obtained (corresponding to the individuals in Supplementary Table 3).

We were only able to compare three males and four females across years to examine changes in syllable repertoire over time at the individual level (Table 3; see Supplementary Table 4 for the number of song selections). Syllable similarity between years varied from 27.3 to 100% for females and from 23.1 to 68.8% for males. There tended to be more syllable deletions than additions (Table 3), but this may be due to rare syllable types going undetected in years with fewer recordings (Supplementary Table 4) where we may have not captured the full repertoire. For the male (1) and female (1) with the most song selections (where we likely captured most of their syllable repertoires), their syllable similarity between two years was moderately high (68.8% and 86.6%, respectively).

## Discussion

Male bellbirds had a larger observed syllable repertoire than females at both the population and individual levels. The population and individual syllable repertoires were on average almost twice the size in males than in females. There was considerable individual variation in syllable repertoire size, ranging from 15 to 32 in males and ranging from six to 16 in females. Such individual differences in repertoire size can arise for a variety of reasons, for example, differences in nutritional stress during early development (Nowicki et al., 1998), learning abilities (Nowicki et al., 2002), density of the local population (Doutrelant et al., 2000), and the health (Buchanan et al., 1999) and dominance of the individual (Voigt et al., 2007). We found more syllable types each year that were sung exclusively by each sex and a smaller proportion of shared syllable types between the sexes compared to Brunton and Li (2006); however, we acknowledge this could be in part due to differences in syllable labelling. Over half of the syllable families were shared between the sexes, but there were more syllable families unique to males than females, with females having only one unique syllable family. The number of syllable types added, deleted and shared between years was similar for both sexes (except for year 1, 2012–2013, where the female syllable repertoire was not fully captured). Population syllable repertoire turnover between years was similar regardless of which two years were compared for each sex, shown by similarity indices, suggesting that syllable types may vary in relative abundance between years rather than being completely lost. This was supported by our finding that syllable types vary in their prevalence within the population for both sexes. Cross-sectional studies tend to be less informative than longitudinal studies for assessing changes in repertoire over time (Gil et al., 2001). However, our analyses of a smaller number of individuals (longitudinal component) produced similar results as to the population analyses (cross-sectional component). For example, both sexes had similar annual changes in syllable repertoires and most syllable types were sung by only a small number of individuals.

To date, few studies have identified significant sexual dimorphism in song repertoires for species where both sexes sing complex, highly variable song. The two song types that female red-winged blackbirds sing are different in structure compared to male song (Beletsky, 1983; Catchpole and Slater, 2008), but there is no detail on what syllable types they may share. Some species have similar syllable structure between the sexes but have differences in the overall or particular spectral parameters of their song (Dutour and Ridley, 2020; Hahn et al., 2013; Wilkins et al., 2020; Yamaguchi, 1998). For example, northern cardinal (*Cardinalis cardinalis*) female song has been described as a neotenic version of male song, but it also differs from males in the amplitude of harmonics and syllable stereotypy (Yamaguchi, 1998). Rufous-and-white wrens (*Thryophilus rufalbus*) also have song types that are unique to males and females, with 18 out of 110 song types shared between the sexes over an 11-year study (Graham et al., 2021), whereas bellbird sexes share no song types (Brunton and Li, 2006). A novel finding of our study is that male and female bellbirds also share very few syllable types – a smaller proportion than reported by Brunton and Li (2006). Male and female bellbird syllable types also differ in spectral parameters, such as fundamental frequency, as found in previous research on bellbird song (Roper et al., 2018; Webb et al., 2021). However, they do share common spectral features at the syllable family level. Sharing such features could be due to limits posed by their vocal organ, the syrinx (Suthers and Zollinger, 2004), or for species recognition (Catchpole and Slater, 2008), as their close relative and competitor, the tūī (*Prosthemadera novaeseelandiae*), also sings complex song (Hill, 2011; Hill et al., 2017). As the sexual dimorphism in song described in our study is so striking and complex, it is unlikely to be solely for sex recognition. Only small changes in the frequency of the ‘fee’ syllable song of black-capped chickadees (*Poecile atricapillus*), for example, appears to be both a signal of sex and individual identity (Hahn et al., 2013). Bellbirds are highly territorial with strong dominance hierarchies (Craig and Douglas, 1984, 1986), and females tend to be excluded from food resources more often than males (Roper, 2012). This could potentially impact their song sharing with males, as females may not have ample learning opportunities to learn song from males if excluded, or there is a cost to singing male-like song when females are already smaller and submissive. With no duetting described between the sexes of this species (Roper and Brunton, personal observation) and female bellbirds singing more in response to neighbouring females than unfamiliar females (Brunton et al., 2008), we suggest females’ unique repertoire functions more in female intra-specific interactions. Understanding whether such costs or benefits to the amount of song sharing between the sexes is occurring needs further research.

Sex-based differences in the bellbird syllable repertoires highlights future directions for studying the functions of bellbird song. Deducing from trends in current literature (Catchpole and Slater, 2008), male song is more likely to be under both inter- and intra-sexual selection (i.e., female-mate choice and male-male competition). Perhaps male bellbirds have more unique syllable types and families than females due to their more diverse functions, such as the ‘sexy syllables’ found in male canaries (Vallet et al., 1998; Vallet and Kreutzer, 1995). On the other hand, if female song is under social selection only (i.e., intra-sexual competition for male parental care and resources, such as food), there may be less pressure to develop a larger repertoire size and unique female syllable types. We also found that there was a greater difference in the syllable repertoire size at the individual level than the population level. This may be explained by the individual variation in syllable families sung by females. For example, we only found one female singing a syllable type from the ‘waah’ family and one female singing a syllable from the ‘pipe-down-squeak’ family, suggesting that there are individual females that learn and sing more ‘male-like’ syllable family types, leading to questions on whether female bellbirds can learn from male tutors. European starling females also have considerable levels of individual variation in repertoire size, with some individuals also singing particular phrase type families that in general are sung by all males (Pavlova et al., 2005). Pavlova et al. (2005) suggested these phrase types may be sung in different social conditions or at different population densities, as an earlier study did not find any females singing these phrase types (Hausberger et al., 1995a; Hausberger et al., 1995b). Hence, the different selection pressures acting on song repertoires need further research to tease apart the functions of distinctive male and female repertoires.

Songbirds can change their song repertoires from year to year (Catchpole and Slater, 2008). At the individual level, we did not capture the full repertoire per individual across multiple years, so we could not address the question of how individual syllable repertoires changed between years. However, for all individuals with sufficient recordings, there were small changes in the syllable types sung between years, regardless of sex. At the population level (cross-sectional analysis), we had the full syllable repertoire for each year (except for females in year 1, 2012– 2013) and the difference in syllable repertoires found between each year was within a similar range. This difference did not depend on whether the comparisons were one year or three years apart (i.e., did not vary with temporal distance). This suggests that bellbirds may be changing the relative abundance at which they sing certain syllable types between years (e.g., a syllable type was rarely sung in one year but common in another). There is potential support for this finding in that individuals either commonly or abundantly sang less than 15% of all syllable types and most syllable types were only sung by a few individuals (a trend found at both the population and individual level). This pattern is similar to one found in Darwin’s medium ground finches (*Geospiza fortis*) where males appear to prefer singing rare syllable types, hence syllable types may change in abundance over the years, rather than being completely deleted from the population syllable repertoire (Gibbs, 1990). In this case, males singing rare syllable types may improve their reproductive success (Gibbs, 1990; Lemon et al., 1992). Hence, for the bellbird syllable types that were considered as either added or deleted between years, some at least might not have been detected each year due to their low prevalence in the population. Microgeographic variation may also explain why most syllable types were sung by few individuals, as in other songbirds this arises through individuals tending to share more syllable or song types with territory neighbours than individuals on distant territories (Barišić et al., 2017; Briefer et al., 2008). However, further studies will be needed to test this hypothesis in bellbirds.

The distinctive level of sexual dimorphism in song repertoires and changes over time have several implications for understanding how each sex learns their song repertoire. It has not been tested whether bellbirds are open- or closed-ended learners. However, we suspect they are open-ended learners based on their high levels of dispersal between populations, but little sharing of syllable types between these populations (Webb et al., 2021). We found that individual changes in syllable repertoires between years was similar to the changes found at the population level. This suggests that the change in population syllable repertoire over time is in part due to small changes in individual repertoires between years (a form of selective attrition), versus juvenile recruitment of novel syllable types or immigrating individuals introducing new syllable types not present in this population. However, we cannot exclude the potential that bellbirds are open-ended learners and that they are learning existing syllable tpes within the population as a contributing factor to this change over time. Our finding that the sexes share a small proportion of syllable types suggests that the individuals may learn, at least to a small extent, from individuals of the opposite sex. Previous research has found that juvenile male and female bellbird songs are more similar to each other in spectral properties than between adult males and females (Roper et al., 2018), also supporting the potential for bellbirds to learn from tutors of the opposite sex, as has been found in a few species (Evans and Kleindorfer, 2016; Geberzahn and Gahr, 2013; Yamaguchi, 2001). Research on tutor choice and learning mode in the species will hence uncover how this species develop their sex-specific song repertoires.

## Acknowledgements

We thank Bridget Hudepol-Maste, Karen Schors, Charlotte Jense, and Xandra Sleeuenhoek for their assistance with fieldwork, and to the University of Applied Science in Agriculture, Food Technology, Environmental and Animal Sciences, Leeuwarden, for facilitating their internship in New Zealand. We thank Chris Smith, Donal Smith, Mhairi McCready, Rachel Shepherd, Victoria Franks, and other Tiritiri Matangi Island volunteers for their help in the field as well. We also express our deep gratitude to the DOC rangers present on Tiritiri Matangi Island during this study. We give great thanks to Barbara Evans for assisting with collecting and processing the sound data. We would like to thank Robert Lachlan for correspondence on using the Luscinia software. Logistical support was provided by the School of Natural and Computational Sciences of Massey University, the Supporters of Tiritiri Matangi Island and the Department of Conservation.

## Author contributions

Michelle M. Roper contributed to the design of the study, developed the methodology, conducted the fieldwork, managed field assistants, processed recordings, contributed to syllable labelling, performed analyses and wrote the manuscript. Wesley H. Webb imported song selections into the software Luscinia and Koe, and labelled syllables. Christine Evans contributed to syllable labelling. Yukio Fukuzawa developed the new software Koe with Wesley Webb, imported syllable selections into Koe, and performed machine learning algorithms for verifying syllable labels. Aaron M. T. Harmer contributed to methodology and assisted with writing the manuscript. Dianne H. Brunton was awarded the funding, designed the study, developed methodology, assisted with fieldwork, assisted with analyses and contributed to the writing of the manuscript. All authors contributed valuable feedback to the writing of the manuscript.

