## Supplementary Methods 1 for "Sexual and temporal variation in New Zealand bellbird song repertoires"

### Supplementary Material

#### Supplementary Methods 1

To verify that our syllable labelling was consistent, two investigations were performed using machine learning. The database of syllables first had to be transformed into their feature representation using 25 different acoustic and spectral features (the list of features can be found on www.koe.io.ac.nz and https://github.com/fzyukio/koe/wiki). The results have different lengths, which correspond to the original durations of the syllables, e.g., the MFCC (Mel-frequency cepstral coefficient) representation of a syllable that is 500ms long is a vector twice as long as that of a 250ms long syllable. The results were aggregated by extracting their means, standard deviations, medians and their dynamic time warp (DTW) distances to several template syllables. After this step, each syllable was represented by a 735-dimensional vector and the t-distributed Stochastic Neighbour Embedding (t-SNE) algorithm was used to assess how well the syllable labels clustered. Then recognition accuracy was assessed using various machine learning algorithms. Supervised machine learning was used to make the computer "learn" how to associate feature values with the actual labels. The trained model could then be used to recognise new instances that the model had never seen before. The recognition accuracy therefore depends on having an appropriate feature representation of the syllables and the correctness of the given labels. High accuracy could only be achieved if both conditions were satisfied. Thus, several different machine learning algorithms (including Gaussian Naïve Bayes, Random Forest classifier, Support Vector Machine, LDA and PCA classifiers) were trained using k-fold validation and their accuracies recorded.
