## Supplementary Figure 1 for "Sexual and temporal variation in New Zealand bellbird song repertoires"

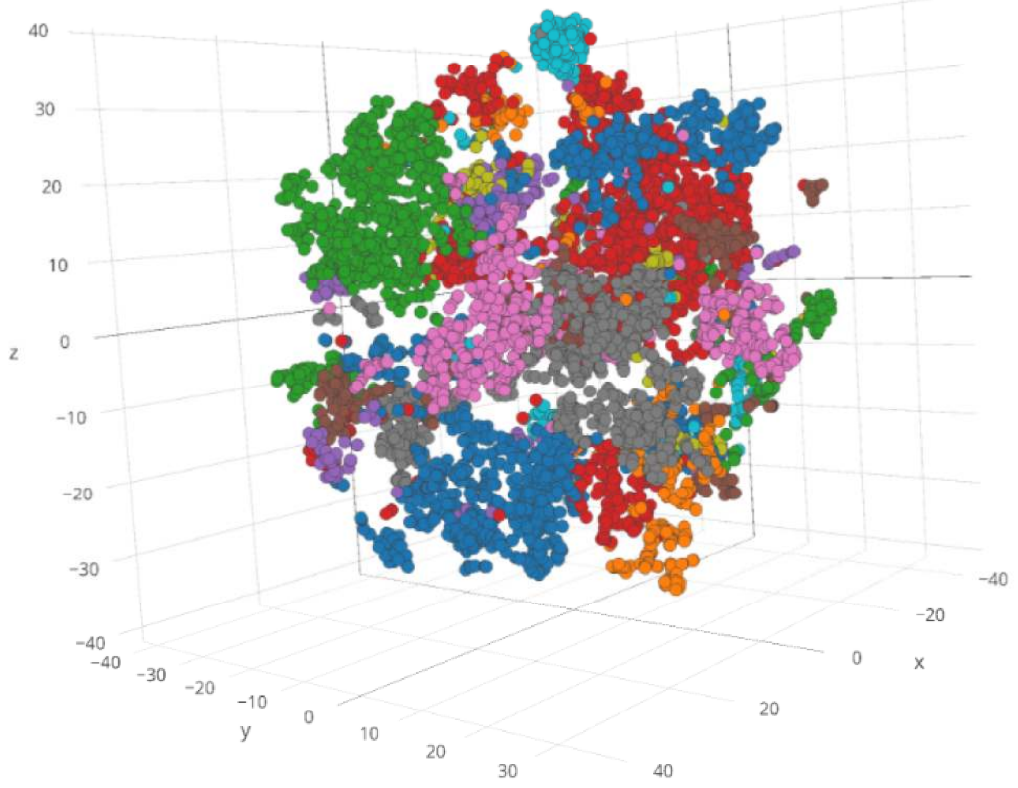


**Supplementary Figure 1.** 3D plot of unsupervised clustering of 7003 syllables using t-SNE (t-distributed Stochastic Neighbour Embedding). Each colour represents the different given syllable labels (although due to there being many label names, the colours are repeated for different labels). Similar syllables cluster together, and the tight clusters of similar colours confirms the labelling of these syllable types.
