## Supplementary Figure 2 for "Sexual and temporal variation in New Zealand bellbird song repertoires"

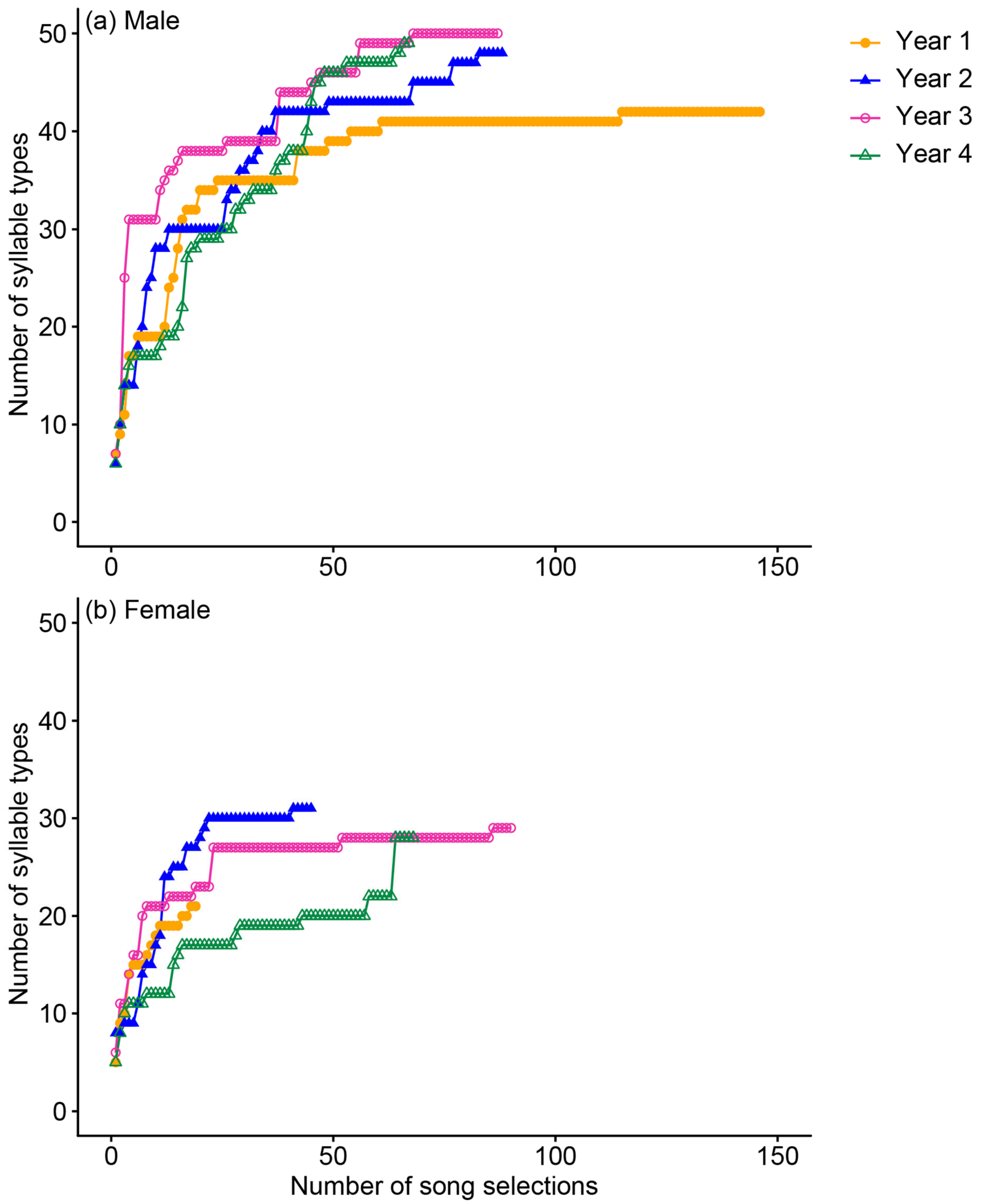


**Supplementary Figure 2**. Cumulative plot of number of syllable types versus number of song selections for male and female bellbirds on Tiritiri Matangi Island for year 1 (2012–2013), year 2 (2013–2014), year 3 (2014–2015) and year 4 (2015–2016).
