## Supplementary Table 1 for "Sexual and temporal variation in New Zealand bellbird song repertoires"

**Supplementary Table 1.** Recognition accuracies of different supervised machine learning methods using k-fold validation (k=25).

| Classifier | Accuracy (%) |
| --- | --- |
| Gaussian Naïve Bayes | 89.5 ± 1.7 |
| Linear Discriminant Analysis | 96.4 ±1 |
| Random Forest Classifier | 82.6 ± 2.26 |
| Support Vector Machine | 96.6 ± 1 |

Note: Each algorithm ran 100 iterations and randomisation was applied at each validation. The result is the mean ± standard deviation of the accuracies.
