## Supplementary Table 2 for "Sexual and temporal variation in New Zealand bellbird song repertoires"

**Supplementary Table 2.** Table of syllable family types with examples of different syllable types (spectrograms with frequency (kHz) on the y-axis and duration (ms) on the x-axis) within each syllable family.

| Syllable family | Spectrogram examples |
|---|---|
| Alarmy | 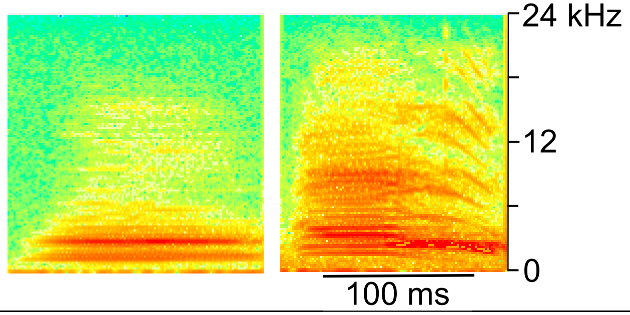  |
| Chortle | 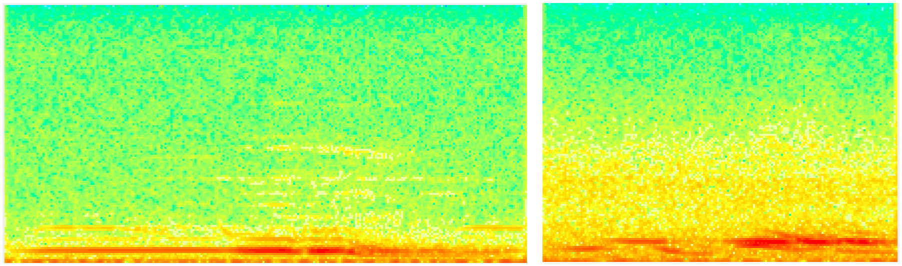 |
| Chump | 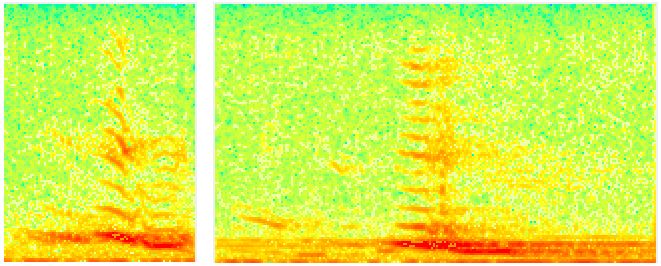 |
| Click | 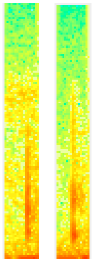 |

| | |
|---|---|
| Cough | 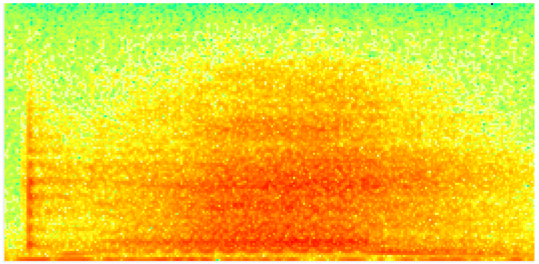 |
| Down-squeak | 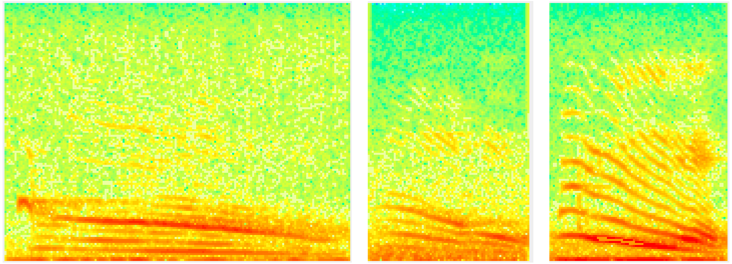 |
| Down-squeak pipe | 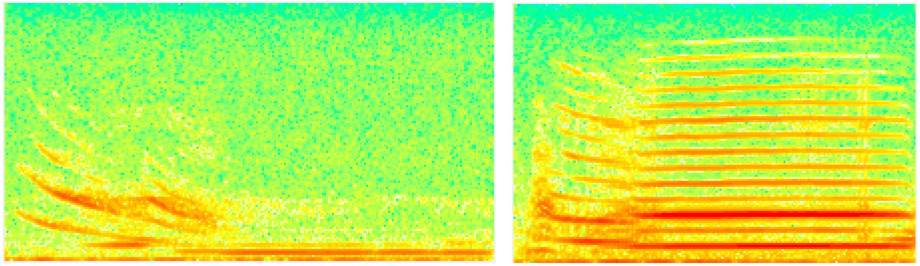 |
| Down transition | 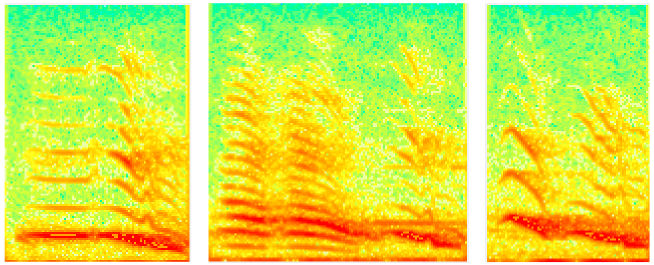 |

| | |
|---|---|
| Flat-squeak | 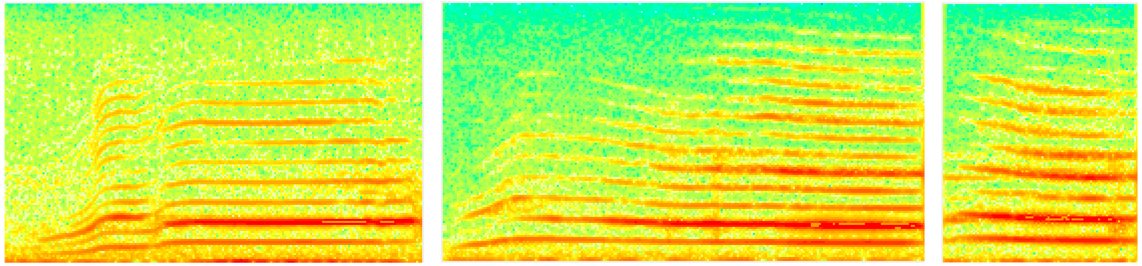 |
| Peak-squeak | 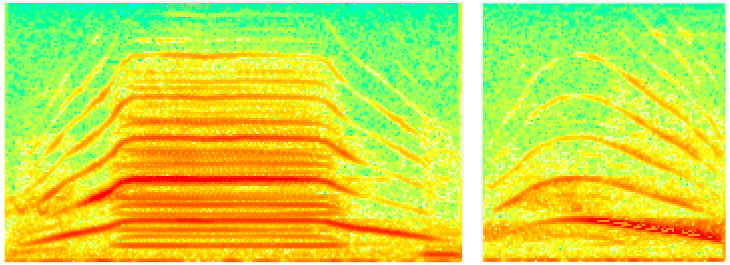 |
| Pipe | 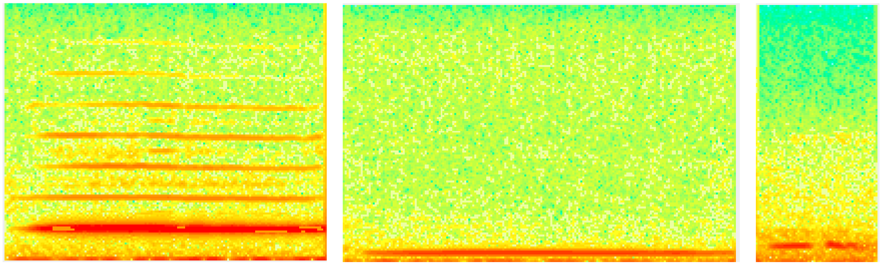 |
| Pipe down-squeak | 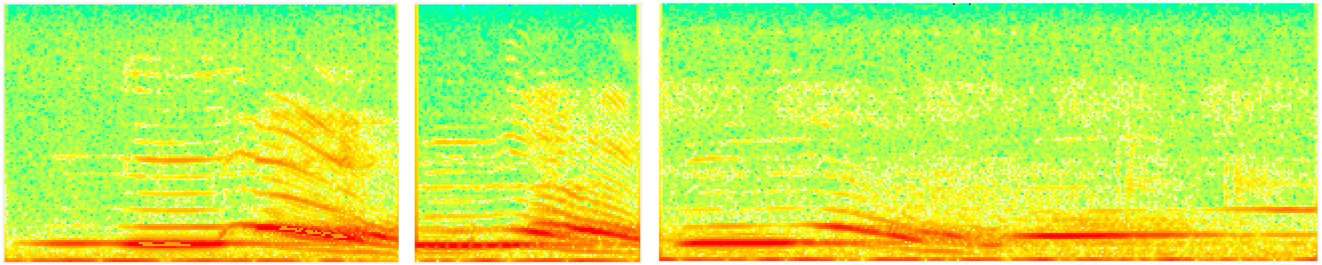 |

| | |
|---|---|
| Step-down | 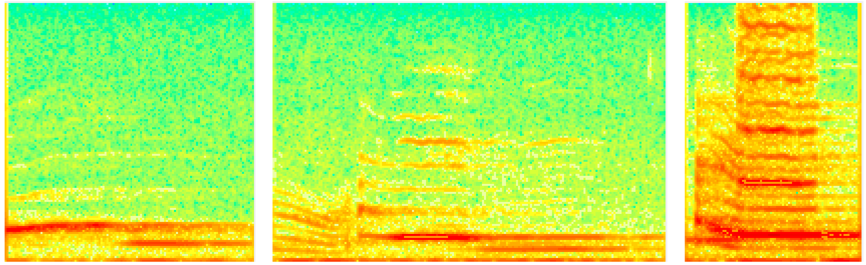 |
| Step-up | 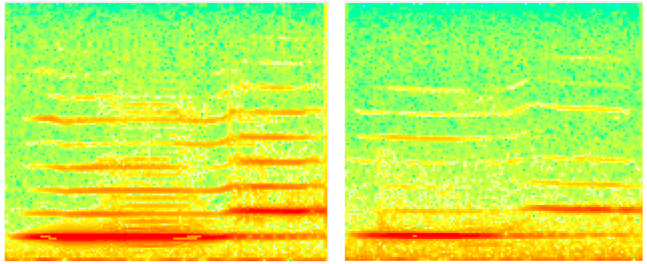 |
| Stutter | 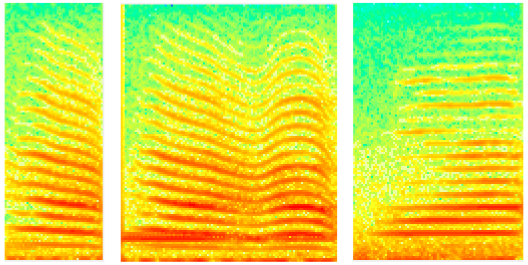 |
| Stutter up-squeak | 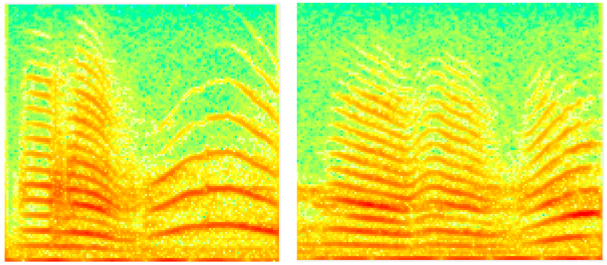 |

| | |
|---|---|
| Trill | 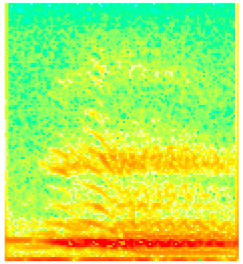 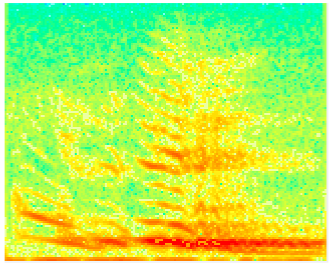 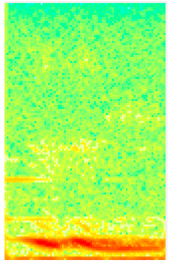 |
| Up-squeak | 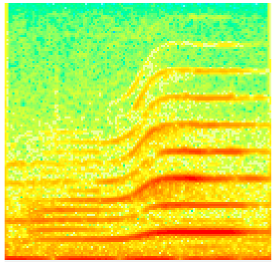 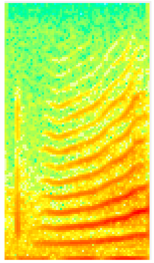 |
| Waah | 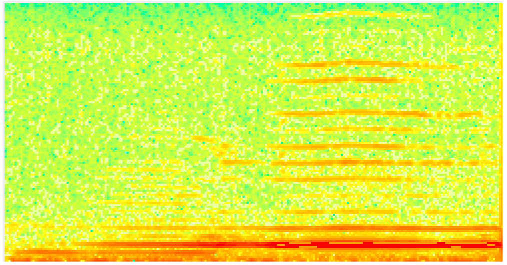 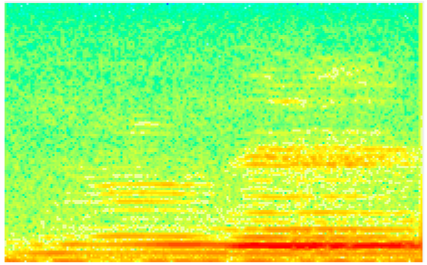 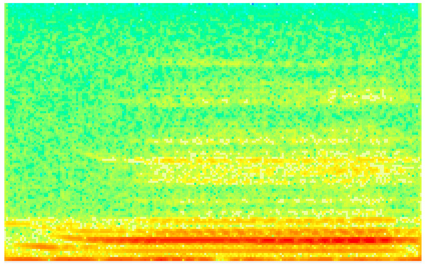 |
| Warble | 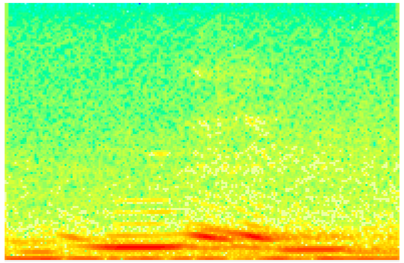 |
