## Supplementary Table 3 for "Sexual and temporal variation in New Zealand bellbird song repertoires"

**Supplementary Table 3.** Syllable repertoire size within each syllable family for male and female individuals with data on number of song selections, recording duration and the time over which each individual was recorded.

| Family | Male | | | | | | | Female | | | | | | | |
|---|---|---|---|---|---|---|---|---|---|---|---|---|---|---|---|
| | 1 | 2 | 3 | 4 | 5 | 6 | 7 | 1 | 2 | 3 | 4 | 5 | 6 | 7 | 8 |
| Alarmy | 1 | | | | 1 | | | | | | | | | | |
| Chortle | 2 | 1 | 1 | | 2 | | | | | | | | | | |
| Chump | 1 | 1 | 1 | 1 | 1 | 1 | 1 | 1 | | | | | 1 | | |
| Click | 1 | 1 | 1 | | 1 | | 1 | 1 | 1 | | | | 1 | | |
| Cough | 1 | 1 | 1 | 1 | 1 | 1 | 1 | | | | | | | | |
| Down-squeak | 1 | 1 | 1 | 1 | 1 | 1 | 1 | 1 | | | | 1 | 1 | | |
| Down-squeak Pipe | 2 | 1 | 1 | 1 | 1 | 1 | 1 | | | | | | | | |
| Down Transition | | | | | | | | 1 | 1 | 1 | 1 | 2 | 2 | 2 | 3 |
| Flat-squeak | 2 | 1 | 1 | | 1 | 1 | | 1 | | | | 1 | | | |
| Peak-squeak | 2 | 1 | | | | | | | 1 | | | | 1 | | 2 |
| Pipe | 3 | 4 | 1 | 4 | 4 | 3 | 3 | 4 | 2 | 1 | 1 | 2 | 5 | 2 | 2 |
| Pipe Down-squeak | 2 | 1 | | 1 | 1 | | 1 | | 1 | | | | | | |
| Stepdown | | 1 | 1 | | 1 | 1 | | 3 | 2 | 1 | 1 | 1 | 4 | 1 | 1 |
| Step-up | 2 | | | | | | | | | | | | | | |
| Stutter | 2 | 1 | 1 | 1 | 1 | 1 | | 3 | 5 | 3 | 3 | 3 | 1 | 3 | 3 |
| Stutter Up-squeak | | | | | 1 | | | | | | | | | | |
| Trill | 3 | 1 | 1 | | 1 | 1 | 1 | | | | | | | | |
| Up-squeak | 1 | 1 | | | 1 | | | | | | | | | | |
| Waah | 6 | 4 | 3 | 5 | 3 | 3 | 2 | | | | | 1 | | | |
| Warble | | | 1 | | 1 | 1 | 1 | | | | | | | | |
| Syllable repertoire size | 32 | 21 | 15 | 15 | 23 | 15 | 13 | 15 | 13 | 6 | 6 | 11 | 16 | 8 | 11 |
| Song selections | 39 | 13 | 26 | 20 | 31 | 14 | 10 | 22 | 14 | 11 | 13 | 21 | 12 | 14 | 10 |
| Recording duration (mm:ss) | 37:40 | 18:21 | 24:54 | 08:59 | 29:20 | 10:37 | 08:37 | 18:12 | 25:21 | 11:16 | 10:45 | 28:03 | 09:20 | 17:19 | 19:23 |
| Time (months) | 10 | 26 | 1 | 0 | 3 | 2 | 3 | 13 | 3 | 1 | 2 | 18 | 14 | 35 | 14 |
