## Supplementary Table 4 for "Sexual and temporal variation in New Zealand bellbird song repertoires"

**Supplementary Table 4.** Number of song selections per year for each individual that was used to compare changes in repertoire size between years.

| | Individual | Year 1 | Year 2 | Year 3 | Year 4 |
|---|---|---|---|---|---|
| Male | 1 | | | 31 | 8 |
| | 3 | 17 | | 5 | 7 |
| | 8 | 5 | 9 | | |
| Female | 1 | | | 17 | 5 |
| | 2 | 5 | 7 | 9 | |
| | 3 | | | 7 | 6 |
| | 4 | | | 11 | 3 |
